# GYDE: A collaborative drug discovery platform for AI-powered protein design and engineering

**DOI:** 10.64898/2026.03.24.714039

**Authors:** Thomas Down, Mateusz Warowny, April Walker, Luigi D’Ascenzo, Donald Lee, Zhenru Zhou, Shengya Cao, Travis W. Bainbridge, John M. Nicoludis, Seth F. Harris, Kiran Mukhyala

## Abstract

As computational tools and machine learning models for protein sciences continue to advance and proliferate, bench scientists face increasing technical challenges adopting these tools for specific applications such as drug discovery. Here we present GYDE (Guide Your Design and Engineering), an open-source, versatile, and web-based collaboration platform designed to make computational analyses of proteins and antibodies easily accessible to bench scientists. GYDE enables the exploration of sequence-structure-function relationships through a tightly integrated visual interface, offering researchers a comprehensive exploration of protein functional determinants either via real assay data or computational tools. GYDE’s intuitive interface facilitates seamless access to cutting-edge AI models for protein and antibody structure prediction, design, and downstream analyses. The flexible and easy addition of new tools and models is facilitated by the use of the Slivka compute API. The platform supports saved sessions that enable researchers to easily share their findings with other users, fostering a more collaborative scientific community. GYDE is freely available for protein scientists in academia and industry to build drug discovery analytics platforms customized to their needs.

## Introduction

Computational structural biology tools are increasingly playing a vital role in the drug discovery workflow. Deep learning methods, such as AlphaFold^1^, RosettaFold^2^, OpenFold^3,4^, Boltz^5,6^, Chai^7^, are now routinely used to predict novel protein structures, conformational changes that may affect a protein’s function, and interactions between a protein to another protein, nucleic acid, or ligand. Tools like ProteinMPNN^8^ have already proven useful for stabilizing target proteins for better protein production or for structural studies. Generative methods like BindCraft^9^ and RFDiffusion^10^ are helping to design therapeutic peptides, miniproteins and antibodies from scratch. Multiparameter optimization methods are helping us improve the properties of therapeutics. However, the rapid rate at which tools are being developed and their increasing computational complexity presents a formidable challenge to their adoption by computational and non-computational scientists alike.

One major challenge in adopting many tools is that each tool is built differently and thus not designed to work cohesively in the same computational environment. Substantial technical expertise and IT investment are needed to link the tools together for the bench scientists to access these. Some noteworthy platform tools that increase accessibility are Jupyter^11^-based sessions like ColabFold^12^, Galaxy^13^, NCBI e-utilities^14^, EMBI-EBI’s Job Dispatcher^15^, Jalview^16^, etc. Assuming all of these platforms are installed, the second challenge is the learning curve associated with each tool as they were often designed for specific purposes and not for a continuous workflow. The analysis of one tool is stored separately from another tool, making collaborative or successive workflows much more difficult. A solution to the above is to rely on commercial offerings that have end-to-end solutions, but this often comes with non-technical limitations, such as costly licenses, proprietary codebase, and limited customizability that could lag behind the release schedule of new tools. The lack of broad and equitable access to state-of-the-art AI models and computational tools slows application of these tools in drug discovery.

The challenge of using these tools is compounded by a disconnect between the execution of the computational task and the integration of the results in an intuitive scientific analysis environment to promote ideation and collaboration. Web-based systems, like Galaxy^13^, facilitate the execution of individual or chained tool workflows but are genomics focused and lack integrated visualizations functionalities necessary for in-depth protein structure-functional analysis. Standalone tools, like Jalview^16^, remain widely used for sequence analyses and alignment, yet are limited in scope for interrogating structure-function relationships and collaboration. Visualization tools, like PyMOL^17^, emphasize the graphical aspects of structural analysis and are more limited for the integration of diverse methods. The sequence-structure-function relationship that originated with Anfinsen’s experiments^18^ is a familiar framework that provides a synchronous and global view of the data that are critical to improving drug molecules. This paradigm also aids in contextualizing computational tools according to their use. AlphaFold converts sequences to structure; ProteinMPNN takes structure and predicts sequences that may improve function; RFDiffusion converts function into a structure. Despite the natural and meaningful significance of this relationship, no tools capture this relationship in an integrated analysis environment.

This manuscript presents a unified and flexible computational platform that integrates these facets to promote efficient collaboration and innovation in protein sciences. GYDE is a web-based platform that not only reduces the barrier to using computational tools but also provides a unified interface allowing for integrated downstream analysis (see Figure 1). This accessibility and democratization fosters interdisciplinary collaboration, enabling scientists from diverse backgrounds to contribute their expertise and perspectives, ultimately enriching the research process. Moreover, it reduces unnecessary complexity and delays in making informed decisions based on models and data, accelerating the pace of scientific advancement as new, transformational methods become available. Herein we describe the GYDE platform and demonstrate its use in six case studies within early-stage drug discovery.

**Figure 1.**
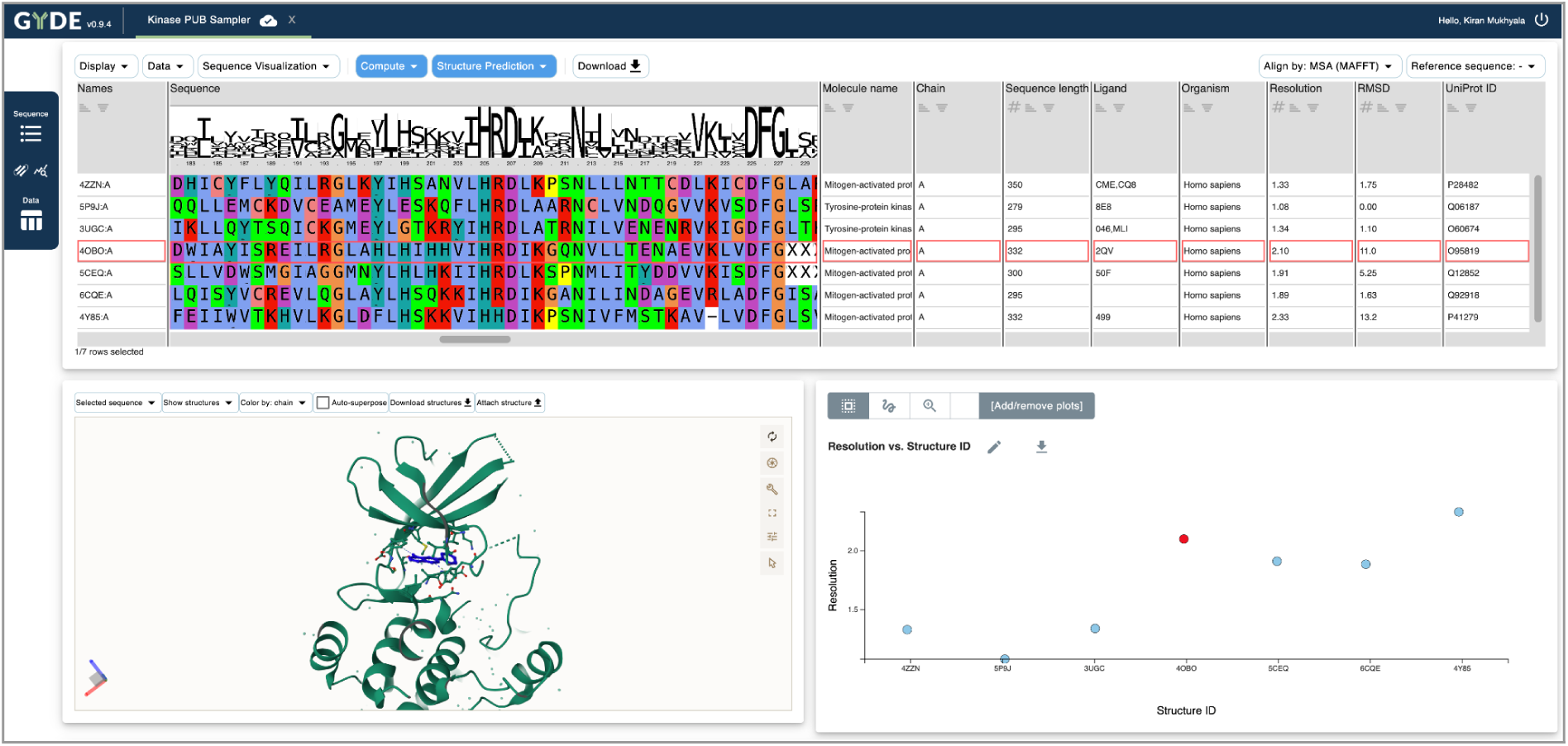
GYDE layout showing the integrated Multiple Sequence Alignment (MSA), Structure Viewer and Plotting panel. The selected sequence in the MSA viewer is used for structure visualization and highlighted in the plotting panel.

## Results

### GYDE Design Principles

In building GYDE, we focused on a set of principles that we felt would best enable protein scientists to use existing and emerging computational tools in their research:

1. A no-code user interface: Unlike solutions that rely on end users to know coding, we felt that a no-code user interface was important for increasing user adoption.
2. Tight integration of sequence-structure-function relationships: Protein scientists are trained to think about this relationship, and we designed an interface that allows integration of the three to enable robust analysis in a single environment. GYDE uses a tabular view that synchronously shows sequence information, corresponding assay data, an embedded structure viewer, and data plots.
3. Access to the latest tools and data: The computational structural biology field is in a stage of rapid acceleration. We knew we needed a backend service runner that was flexible and allowed new tools to be added easily for efficient evaluation and timely usage. Additionally, we integrated key existing public data repositories (e.g. PDB, UniProt) to access relevant contextual knowledge.
4. Collaboration: The sharing of structure, sequence, and functional data can be a barrier to collaboration as they require different tools (spreadsheets, sequence viewers, structure viewers) and files need to be shared individually. By storing user sessions in integrated datasets, we enable robust data storage and sharing with collaborators via an intuitive hyperlink sharing feature.

### System Architecture

GYDE is built using a modular architecture to help isolate the development of components for user interface, computation, and data management, while still allowing of inter-module communication (see Figure 2).

**Figure 2.**
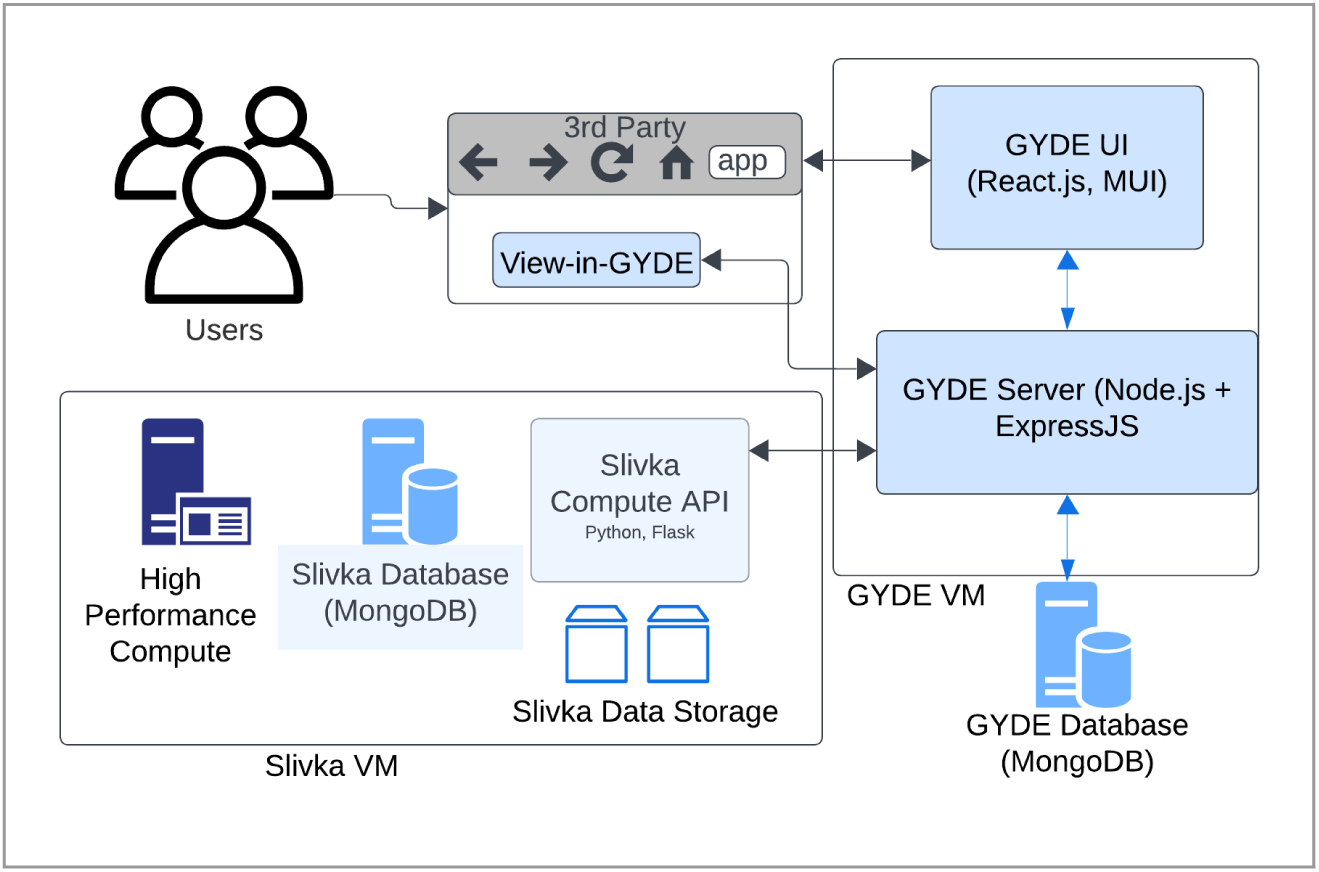
GYDE architectural components showing the integration between GYDE frontend, backend, databases, Slivka and third-party integration.

#### User Interface and Interaction

At the heart of the GYDE architecture is the web interface (GYDE UI), which is organized into distinct but integrated components. This provides flexibility for each user to manipulate the components for an intuitive and seamless experience. Commonly used components are purposefully made more accessible by default, as described below.

The main analysis page consists of the following visual analysis components:

1. Multiple Sequence Alignment (MSA) Viewer: Many workflows start by reviewing the protein sequences. The sequences can be aligned using MAFFT^19^ for proteins, or via an antibody-specific aligner such as Absolve^20^ that annotates the complementary-determining regions (CDRs). The MSA Viewer was built as a tabular format as this allows for further customization. Some examples include filtering, sorting, selecting, annotating, commenting, and highlighting sequences.
2. Structural Visualization: The Mol* Viewer^21^ structure visualization component enables users to explore molecular structures associated with one or more sequences selected in the MSA viewer. This sequence-structure integration between the MSA and Mol* viewers is achieved by automatically standardizing the residue numbering between a sequence and structure via sequence alignments. Running structure prediction methods like Alphafold^1^, Chai-1^7^, Boltz-1^5^ or ABodyBuilder^22^ automatically adds the results to the viewer with optional superposition. Users can manipulate the view, toggle different structural features, and perform structure-based analyses using Mol*’s built-in capabilities. This also serves as a portal whereby users can upload additional or download predicted structure models from the GYDE-embedded tools.
3. Plotting: The plotting component supports the rapid creation of histograms or scatter plots based on the data values uploaded into the GYDE data table. These data values typically represent the functional properties of the protein or mutants being analyzed. Interactive selection of data points in the plotting module automatically selects the corresponding sequences in the MSA module thus allowing a powerful multi-axis exploration of the sequence or structure-function relationship.
4. Sequence-To-Image Viewer: Many computational analyses produce summaries in the form of static images. For example, structure prediction tools, like Alphafold, produce plots of MSA coverage and confidence metrics. Wet lab experiments often include images of their results for example, sensorgrams from Surface Plasma Resonance (SPR) experiments. Datasets that include such images for one or more of the protein entries can be viewed in GYDE for comparative analysis and reasoning of results.
5. Frequency Analysis: A common task in analysis of sequence designs is identification of conserved residues or filtering sequences with certain residues. The amino acid Frequency Analysis component aids such tasks by showing distributions of amino acids in selected positions or those with selected conservation levels (Figure 6).
6. Heatmap viewer: Both modern experimental and computational datasets can produce complex, rich datasets with saturating mutagenesis at each sequence position. GYDE provides a heatmap view to provide concise and interactive visualization of this data matrix, which aids navigating and distilling such information (Figure 5).
7. Sequence logo: GYDE also provides an integrated sequence logo viewer, obviating the need for users to seek other software for this common and informative visualization method for protein variability by position.

#### Computational Integration with GYDE Server

The GYDE Server acts as a central hub that orchestrates the system’s computation and data management operations. It handles requests to send and receive data from GYDE UI. To run computational tools, GYDE integrates with the Slivka Compute API^23^, which provides job runners for various high-performance computational resources such as LSF, Slurm, Sungrid engine, or local computing resources. This integration with Slivka allows GYDE to leverage external computational power for large-scale data processing and analysis, thereby enhancing GYDE’s capabilities and scalability. Table 1 provides a list of Slivka services currently available in our in-house deployment of GYDE.

**Table 1.** Sample list of tools available in an internal deployment of GYDE, provided by Slivka or other internal servers.

| Slivka Service | Analysis Description | Model type |
| --- | --- | --- |
| MAFFT<sup>19</sup> | Multiple Sequence Alignment | Alignment |
| AlphaFold2<sup>1</sup> | Structure Prediction | ML |
| Boltz1/2<sup>6</sup> | Structure Prediction | ML |
| Chai-1r<sup>4</sup> | Structure Prediction | ML |
| OpenFold3<sup>4</sup> | Structure Prediction | ML |
| MOE Ab_workflow<sup>24</sup> | Structure Prediction | Physics-based |
| ABodyBuilder2<sup>22</sup> | Structure Prediction | ML |
| Ibex<sup>25</sup> | Structure Prediction | ML |
| Therapeutic Antibody Profiler<sup>26</sup> (TAP) | Ab surface properties | Physics-based |
| MolDesk<sup>27</sup> | Ab surface properties | Physics-based |
| Absolve<sup>20</sup> | Ab Fv numbering | Alignment |
| RaSP<sup>28</sup> | Protein stability | ML |
| Rosetta $\Delta\Delta G$<sup>29</sup> | Protein stability - Free energy of folding | Physics-based |
| ProteinMPNN<sup>8</sup> | Inverse Folding - protein sequence design | ML |
| ThermoMPNN<sup>30</sup> | Inverse Folding - protein stability | ML |
| LigandMPNN<sup>31</sup> | Inverse Folding - ligand-conditioned protein sequence design | ML |
| BindCraft<sup>9</sup> | De novo protein design | ML |

#### Data Model and Management

The foundation of GYDE’s data management system is the dataset, which functions as the core unit for storing and sharing information within a centralized repository with access control. This structure supports real-time collaboration, allowing multiple researchers to work concurrently on shared projects.

A GYDE dataset is designed around a flexible, columnar dataframe integrated with key metadata attributes. In this arrangement, rows correspond to individual molecular assemblies (such as monomers or complexes), while columns store corresponding experimental measurements or computed values. This model is highly adaptable, supporting specialized, domain-specific data types critical to drug discovery, including protein, DNA, and RNA sequences, as well as cheminformatics formats like SMILES. The design also incorporates hidden columns to efficiently cache computational results and maintain stable identifiers.

To ensure robust persistence, scalability, and seamless sharing of analysis context, GYDE uses a persistent core data model backend. This system enables complex operations, such as filtering data and merging datasets, efficiently even with large volumes of information. Datasets are secured with user identifiers and access control flags, supporting private, shared, and public modes. Furthermore, versioning protocols are in place to accommodate schema evolution, ensuring the system remains adaptive to future changes in the data landscape.

Integration with both internal and external data is a key feature. GYDE connects directly to major public protein databases, including PDB, UniProt, and Pfam, allowing users to instantly retrieve contextual structural and sequence data. It also integrates with proprietary laboratory information management systems (LIMS). A critical component for streamlining workflows is the “View-in-GYDE” API endpoint, which accepts data directly from external applications. This feature bypasses manual export/import steps, significantly enhancing user adoption by facilitating the immediate visualization of sequence-associated data—for instance, merging antibody library data with experimental results like Surface Plasma Resonance (SPR) or protein expression values. The modular design of the architecture ensures flexibility and scalability, making it well-suited for modern data-driven protein science environments.

### Case Studies

GYDE has been applied in several case studies, demonstrating its utility in protein engineering and design projects. Users report significant time savings (in some cases from multiple days down to minutes or hours) and enhanced collaborative efficiency. The interactive visualization capabilities have been particularly appreciated for aiding intuitive understanding of protein sequence-structure-function relationships. Importantly, the platform provides benefits in both standardized, large-scale pipeline work as well as *ad hoc*, individual creative scientific explorations. The following sections present some of these case studies in using GYDE for protein science.

#### Structure Prediction

GYDE facilitates access to a growing list of structure prediction methods, particularly AlphaFold2 and emerging co-folding methods such as Chai, Boltz, and OpenFold. These tools address the common need of modeling complexes and multimer systems not available in public repositories, whether with protein partners or with small molecules, nucleic acids, or modified protein systems.

##### Single-pass TransMembrane (STM) Multimers

We used AlphaFold2-multimer^1^ to generate protein complex predictions for putative interactions between single-pass cell-surface proteins based on an experimental proteomics dataset of 1,381 proposed interactions, with the aim to ascertain whether the computational prediction could provide guidance to inform validity of the broad experimental results and highlight the most promising partnerships and biology for further study. A similar approach has now been documented for study of orphan ligand receptors^32, 33^. Predicted structures were evaluated on their overall confidence score^34^, the pDockQ^35^, and the interaction interface size (via PISA^36^). We removed low confidence regions of the predicted structures. With GYDE’s data upload mechanism we merged these predictions and computational metrics with experimental screening values and public database knowledge on any relevant existing structures of the proteins of interest and whether the pairing is known in databases such as STRING^37^. Plotting the standard deviation of the confidence score versus the maximum confidence score among five models generated per complex allowed us to visually identify distinct populations of our predictions compared to the orthogonal experimental data or literature information on the proteins mapped in the color channel (Figure 3B).

**Figure 3.**
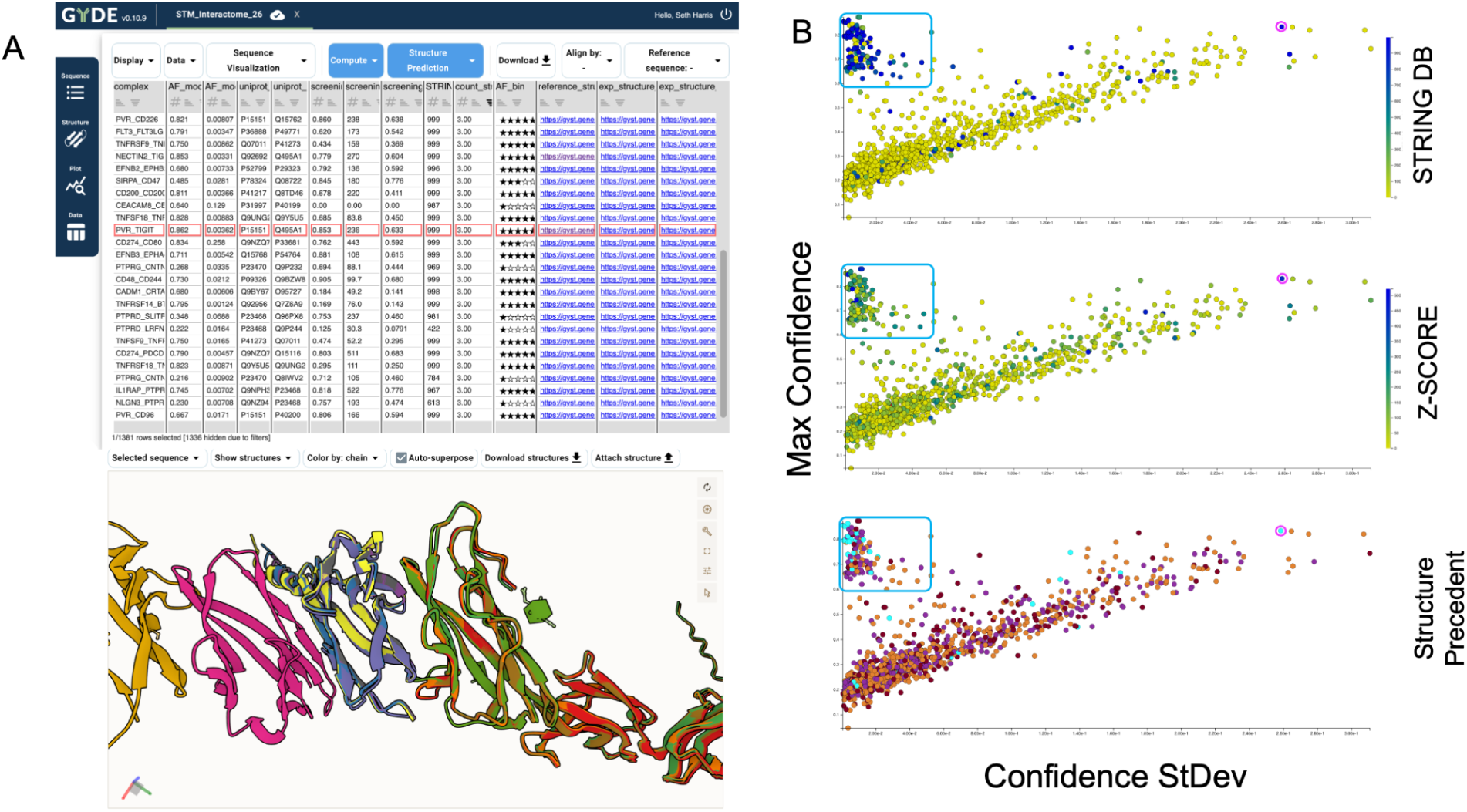
GYDE navigation of the interactome dataset. (A) The GYDE table and structure viewer filtered for partnerships with known experimental complexes. The structure viewer is synchronized to the tabular selected high-confidence row (e.g. PVR:TIGIT) showing an overlay of AlphaFold2 predictions versus the reference (PDB 3UDW). (B) GYDE plots by AlphaFold2-multimer prediction maximum confidence versus confidence standard deviation for 1381 potential protein complexes. The blue box indicates a region of reproducible, high-confidence predictions, while regions to the right indicate more variability across the five predictions per sample, while still indexed to the top score among the five results, such as for the magenta circle highlighting PD-L1(CD274):CD80 as a known interaction example. The top plot is colored by STRING DB score as an indicator of prior knowledge of interaction. The middle plot is colored by the experimental screening Z-score from this work. The lower plot is colored by categorical indication of experimental structure precedent status in the PDB. Cyan: structure(s) with both partners in complex; purple: individual structures for both partners independently; orange: at least one or the other partner has a known structure; dark red: no structures known for either partner.

In this manner we could rapidly distill the large complex dataset using GYDE’s interactive capabilities and user annotation fields to isolate and annotate key categories of results. Of the 1,381 experimentally proposed potential partnerships, only 45 (3%) had prior structure information of the relevant complex, while 402 (29%) of the set had structural information of both components individually, 670 (49%) have a structure for only one of the components, and 264 (19%) have no prior experimental structures of either partner protein. The upper left region of the plot (Figure 3B, blue box) indicates strong confidence predictions, suggesting higher likelihood of interaction and greater promise to determine experimental structures. As expected, this sub-population is enriched for known interactors in STRING database, higher experimental validation, as well as pre-existing structural knowledge (Figure 3B, top). Nonetheless, members of this population not represented in STRING or lacking existing experimental structural knowledge afford opportunities for novel discovery that have a relatively high chance of success. Of further interest, the region in the upper middle and right of these plots indicate cases where some of the predictions scored with confidence, but there was more variability across the 5-prediction set (hence higher standard deviation to the right on the plot). Yet the presence of some known interactors (higher STRINGDB scores), some of which include known structures of the complex (e.g. PD-L1:CD80 marked by the magenta circle in Figure 3B) act as a positive control, showing that despite the inconsistency in structure predictions, these points nonetheless indicate some potential for informing physiological validity. Finally, the extensive number of examples towards the lower left in each plot, indicative of reproducibly poor structure prediction confidence, is notable. While some may be genuine negative results to suggest these are not real interactors, it is undoubtedly true that bona fide interacting partners are also here as false negatives. This highlights challenges in this regime of likely transient and weaker interactions typical of cell surface proteins or glycosylation and other modifications that were not performative in these prediction algorithms and provides a focus for continued attention in model development.

##### A comparison of Boltz-1 vs Chai-1r

Many large scale computations on proteins published previously^38,39^ lack a data exploration tool that’s especially geared towards non-computational users. To exemplify GYDE’s capabilities to navigate complex structural datasets, we re-generated the recent co-folding benchmark dataset Runs-N-Poses^39^ with Chai-1r and Boltz-1 and imported these results using GYDE’s data upload feature. This enabled us to filter and sort results by the prediction metrics and to inspect prediction quality and discrepancies via the integrated Mol* viewer (Figure 4A). We were able to reproduce the findings from the Runs-N-Poses study showing that these methods perform well when the similarity to training data is high, but don’t generalize well to unseen data (Figure 4B).

**Figure 4.**
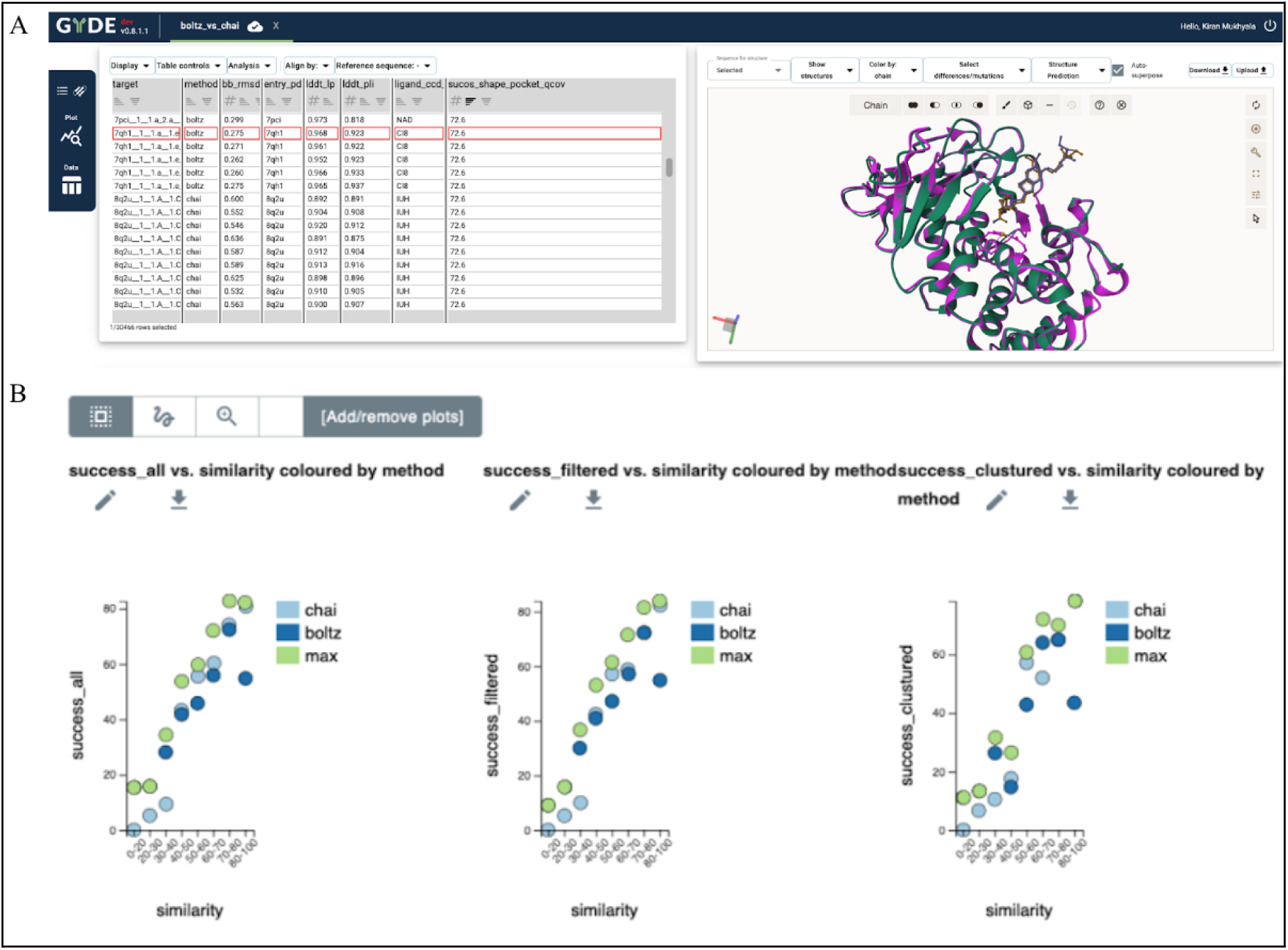
Exploring the results of the Runs_N_Poses dataset re-generated only on Boltz and Chai. (A) GYDE dataset to explore individual predictions showing superposition of predicted vs ground truth co-folding methods. (B) Prediction accuracy vs. training set similarity to validate the Runs_N_Poses results (Fig 1A, B, C from the original publication). Success rate is defined as the percentage of system ligands with <2Å RMSD and >0.8 LDDT-PLI, for ligands in each similarity bin for each method and when selecting the best-scored model from the two methods, for all system ligands in the common subset, only system ligands having <100 training systems containing analogs, and only cluster representatives.

#### Antibody Engineering

##### Enablement of Antibody Engineering Workflows

Antibody engineering often aims to enhance binding, reduce liabilities, or improve developability by modifying the least number of residues from a lead candidate. The workflow involves multiple analytical steps and software tools as described:

1. **Annotation**: Identifying antibody CDRs and framework regions according to various numbering schemes (Kabat, IMGT, Clothia). Requires a germline segment annotation software.
2. **Clonotyping**: Grouping similar antibodies by VGene and CDR3 identities. Requires clustering and filtering tools.
3. **Mutation Analysis**: Portraying somatic mutations within clonotypes with functional data. Requires MSA and table merging tools.
4. **Structure Mapping**: Mapping mutation-function relationships onto antibody structures. Requires a structural modeling software with custom functions.
5. **Residue Modification**: Deciding which residues to modify to achieve objectives. Requires custom tools.

GYDE simplifies the core steps by integrating antibody-specific annotation tools, assay/computed properties, predicted antibody structures, heatmap view of the variants, and a picklist generator. To showcase GYDE’s capabilities to run through a typical antibody engineering workflow, we started with a public antibody. Given the publicly available structural and NGS sequencing data associated with anti SARS-CoV2 antibodies, we arbitrarily chose an antibody called C102^40^ to represent the “lead” candidate that a research group may be interested in. A common next step is to gather data on the antibody mutations in an attempt to approve an intrinsic property, such as binding affinity. This can be done experimentally by performing saturation mutagenesis of the CDR residues and identifying which residue, or residues, improve binding^41^. Another strategy is to mine B-cell repertoire sequences to find similar clonotypes and observe which mutations were selected^42^. Using the latter approach, we mined the OAS database^43^ for SARS-CoV2 patients’ B-cell repertoires^44,45,46^ and extracted sequences for which the heavy chain CDR3 were within 90% identity of the lead antibody. The relevant sequences were then uploaded into GYDE to initiate the workflow as shown in Figure 5. The procedures used to generate this example data are provided in supplementary file Supp_ExAntibodyCoV2.xlsx.

**Figure 5.**
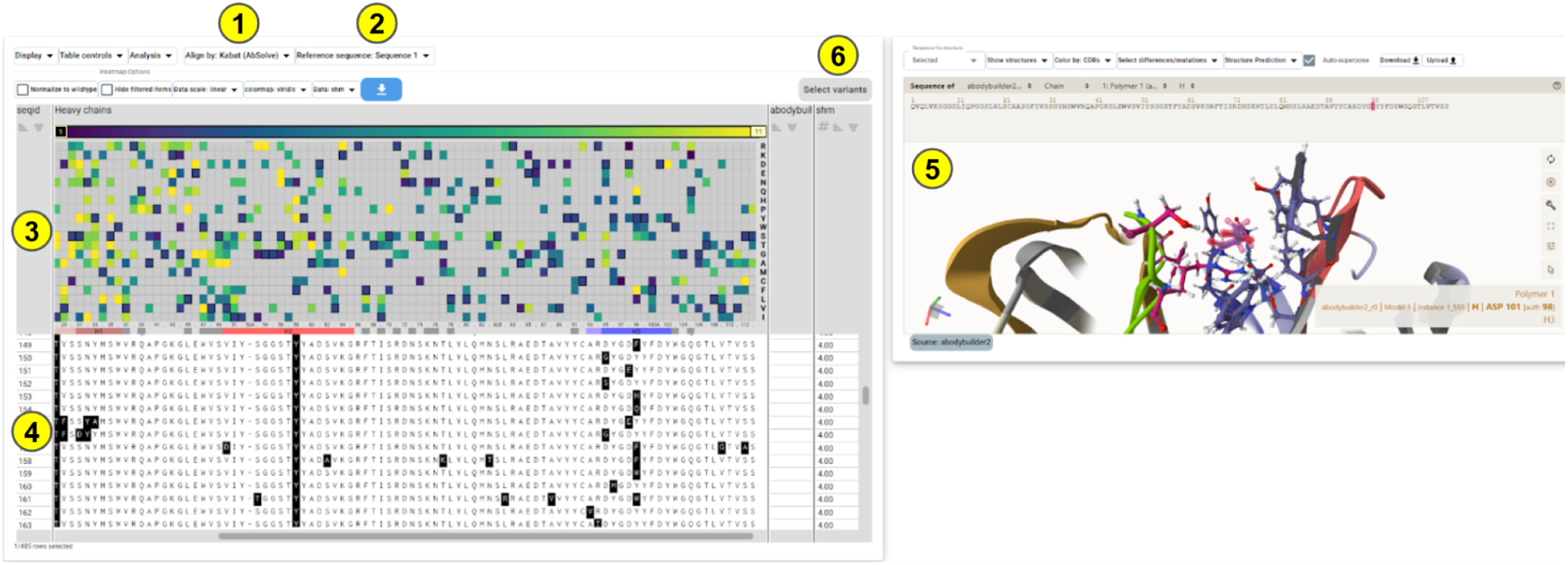
GYDE enables antibody rational design workflows shown here mapped to the graphical interface. 1) set antibody numbering scheme. 2) choose a reference antibody for relative comparisons. 3) view a data column as a heatmap that is aligned to the residue numbers. 4) click a heatmap grid to dive deeper into sequence variabilities. 5) predict the antibody structure to review heatmap-sequence-data relationships. 6) select variants of interest to synthesize.

Using GYDE’s predicted antibody structure and heatmap viewer, the variability in the CDR3 region of the heavy chain readily shows preferences for certain residue properties at specific positions. We inferred that these residues may have experienced a selection pressure, either due to structural constraints or binding preference to an antigen epitope. One position in particular is HCDR3, 98 (Kabat numbering), which is void of positive-charge residues but tolerant of negative-charge Asp and Glu. An obvious inference is that this position may be proximal to a positively charged residue in the Spike receptor binding domain (RBD). The predicted structure by ABodyBuilder2^22^, invoked directly in GYDE, showcases the side chain is likely facing towards the antigen as opposed to being buried inward. By comparing with the experimental complex structure 7K8M^40^ there is indeed a proximal Lys (417) residue in the Spike RBD for which contact is feasible. While this is a retrospective observation given that an experimental structure is available it demonstrates the potential for hypothesis generation. This information provides engineering constraints for improving the antibody properties beyond binding affinity, and therefore having a mechanism to guide the next iteration of design is beneficial. On this note, we directly integrated a picklist generator into GYDE by repurposing the heatmap viewer as a sequence editor, which skips the need for a separate editing tool. Overall, this engineering workflow can be readily applied for other antibodies against different targets, and without involving any code. Therefore we envision that GYDE provides an infrastructure for computer-aided design of antibodies, where computational and protein scientists can interrogate impacts of mutations across sequence, structural, and functional regimes, all done in one collaborative session.

##### Anti-PD-1 antibody

As an example of integrating computationally engineered antibodies with experimental data, we leveraged the GYDE platform to design mutants of an anti-PD-1 rabbit antibody using ProteinMPNN, as detailed in our published work^47^. The study explored the engineering of a human-mouse cross-reactive monoclonal rabbit antibody, h1340.CC, which features a non-canonical disulfide bond that posed a challenge due to its potential impact on antibody developability. Initial attempts at removing the disulfide bond led to a loss in PD-1 affinity, prompting the use of machine learning and structural guidance to develop a new variant with improved properties. ProteinMPNN integrated in GYDE together with the embedded Mol* structure viewer, amino acid frequency analysis, and experimental data-merge capability facilitated the rapid generation and selection of sequence designs, allowing the team to efficiently create a variant that restored affinity while maintaining favorable developability characteristics. As described in the study, the crystal structure of h1340.CC Fab bound to human PD-1 was uploaded to GYDE and the heavy chain was selected in Mol* to use as input to ProteinMPNN. The user specifies the desired number of designs and the temperature parameter. After a few minutes of run time, the ProteinMPNN generated sequence proposals are automatically displayed alongside a Frequency Analysis widget in GYDE that allows the user to see the distribution of amino acids at one or more selected columns in an MSA (Figure 6). Analyzing the co-frequencies of disulfide cysteines (C35-C50), we honed in on the most frequent non-cysteine variants in the proposals, with Ala-Ala (AA) being the most common (22%), followed by Ser-Ala (SA, 11%) and Val-Ala (VA, 9%). Running Rosetta ΔΔG analysis on these variants in GYDE supported experimental verification of higher affinity for the AA variant over the SA variant. Within this subset (AA, SA and VA), we filtered mutations to those with frequency of 50% or higher (58 mutations), and subsequent visual inspection with the Mol* widget to ensure solvent-facing and non-antigen-binding region characteristics (6 mutations). Experimental measurements of affinity for the single-mutation variants with the SA background were then uploaded and merged with the designs, highlighting three of these with improved affinity. These were combined to select the final variant that demonstrated affinity levels comparable to the original CC antibody. This integration of structure- and ML-guided approaches showcases a promising methodology for antibody engineering, emphasizing the potential of GYDE and ProteinMPNN in overcoming similar challenges in therapeutic antibody development.

**Figure 6.**
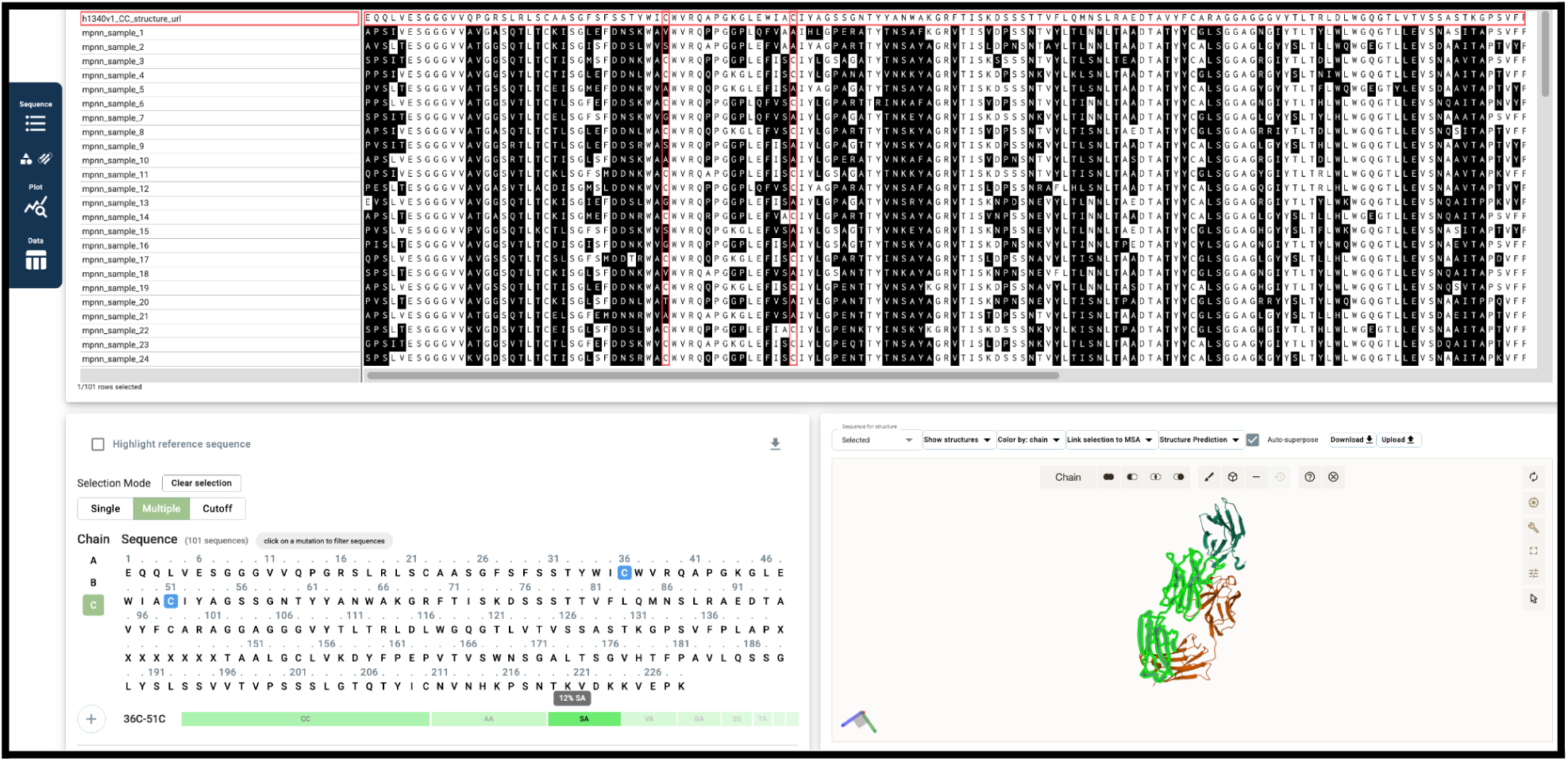
The GYDE interface during ProteinMPNN sequence design of the antib-PD1 antibody heavy chain. The sequence viewer displays the proposed sequences from ProteinMPNN highlighting mutation occurrences. The structure viewer can be used to show the reference structure highlighting design positions or specific columnar selections. The Frequency Widget at lower left provides an interactive workbench to study mutation distributions at each site and make selections manually or based on these frequencies.

#### Protein Design

##### Engineering HyperTEV design with ProteinMPNN in GYDE

Exemplifying GYDE’s efficiency in defining protein design space, we replicated the computational components of a published study yielding improved catalysis from an engineered version of the Tev1 protease^48^. Those authors leveraged ProteinMPNN on various subsets of less conserved positions in the sequence, achieving a ∼20x improvement in catalytic efficiency with the best three of their designs (termed HyperTev60, 56 and 89). Using GYDE, we uploaded a single chain from the PDB structure 1LVM^49^. While the design space can be manually selected on the sequence display, or via the embedded 3D Mol* viewer, definition of complex or extensive data-driven selections such as used here (namely only allowing less-conserved residues to vary and excluding those proximal to the substrate site) is facilitated by a flexible dialog input. This flexibility facilitates a variety of data-driven or conditional selections (ligand or active site proximity, interface involvement, surface exposure, conservation, etc.) to be readily established and confirmed in the linked 3D display (Figure 7A).

The GYDE interface to ProteinMPNN includes settings for key run parameters (Figure 7A). Following the best results in the published study, we applied a 0.2 sampling temperature, 0.2 backbone noise, and generated 100 proposed sequences allowing variations at 91 positions within the defined design space. GYDE’s sequence logo, heat map depictions, and the interactive frequency widget selector readily quantify the complex mutational landscape of the output results (Figure 7B). More than 95% of the design positions included residue proposals found in the top 3 successful HyperTev designs (HyperTev60, 58, 80), demonstrating the GYDE platform can be efficiently used for protein engineering efforts. (Supplemental Figure 2)

**Figure 7.**
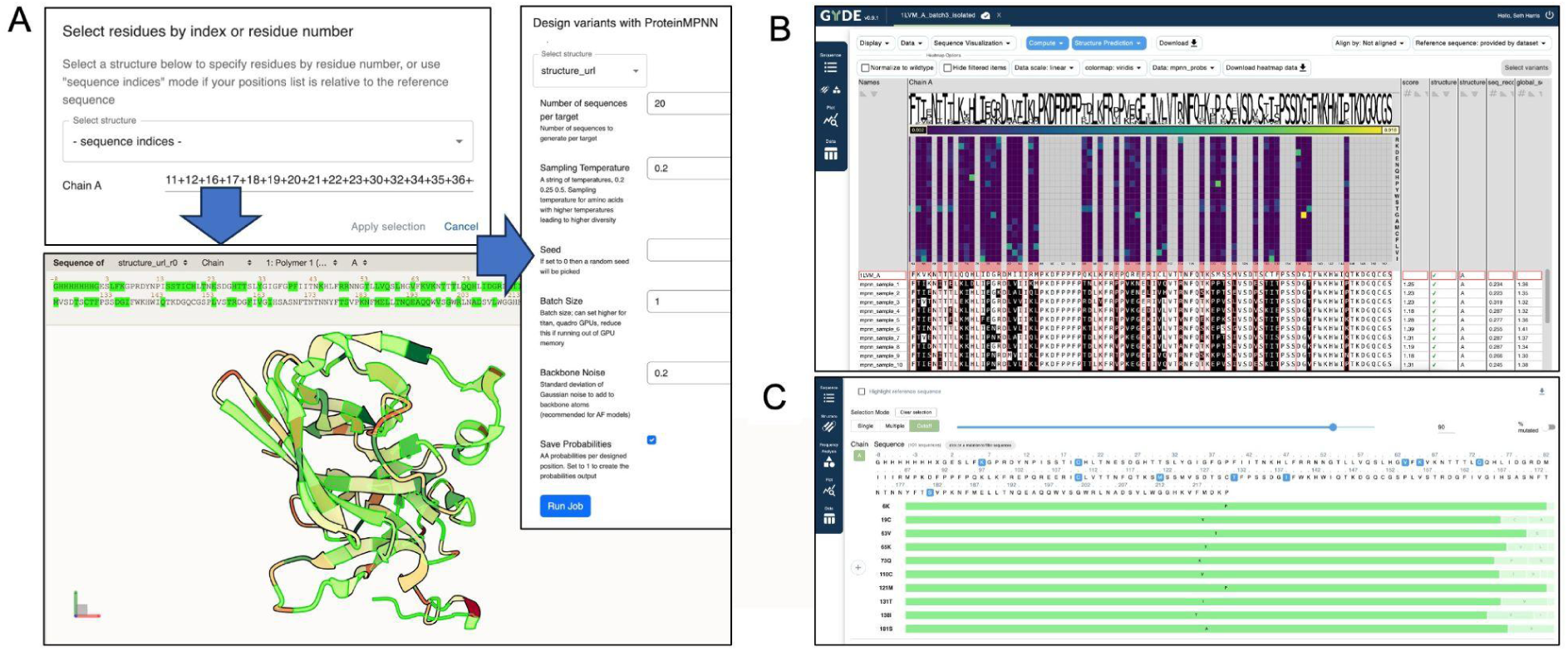
GYDE implementation for a ProteinMPNN design workflow applied to engineering Tev protease. (A) Complex design space selections can be designated by sequence indices, confirmed in the structure viewer and simple dialog options control the algorithm run. (B) The design proposal output is viewed in the multiple sequence alignment including sequence logo, heat map, and sequence-associated meta data (e.g. design scores) and user data that can be used for sorting and filtering results. (C) The frequency widget allows interactive analysis and selection of key mutations and distributions.

##### Design of LRRC15-binding miniproteins in AAV scaffolds

Miniprotein binder design using generative protein models can now routinely provide submicromolar binders to targets of interest ^10,9^. This shows promise for many therapeutic applications, one of which is the incorporation of retargeting elements into AAV capsids (Figure 8A) by incorporating binders into loops in the capsid protein. We used RFDiffusion^10^ with ProteinMPNN^8^ and BindCraft^9^ to demonstrate the successful design of a LRRC15 binder that could be incorporated into AAV capsids. LRRC15 is a membrane-receptor biomarker for cancer-associated fibroblasts and a potential therapeutic target^50^. Binders to LRRC15 could serve as imaging reagents or enable the targeted delivery of therapeutic payloads, as exemplified here.

Using an AlphaFold model of murine LRRC15, we selected five hotspot regions on the mouse LRRC15 extra-cellular domain (Figure 8B) and used these regions as target surfaces for design in BindCraft, RFDiffusion and ProteinMPNN. We uploaded the designs in GYDE and used selection and filtering to narrow down the design set from more than 1000 design candidates down to a testable set of designs. Together, we tested 120 candidates from RFDiffusion and 55 candidates from BindCraft for their LRRC15 binding and overall VLP yields. We merged these experimental data into a GYDE session to analyze the results (Figure 8C,D). We discovered a number of BindCraft designs that had high mLRRC15 binding without decreasing VLP yields (Figure 8D), though overall, more of the BindCraft designs tended negatively to affect VLP yield. On the other hand, RFDiffusion designs were better tolerated as VLP fusions, but none had as strong binding to LRRC15 as the top Bindcraft proposals.

GYDE allowed us to rapidly design binders, share the designs with colleagues, and analyze the results within the same environment including the import of experimental affinities, promoting rapid and collaborative use of protein design tools in drug discovery workflows.

**Figure 8.**
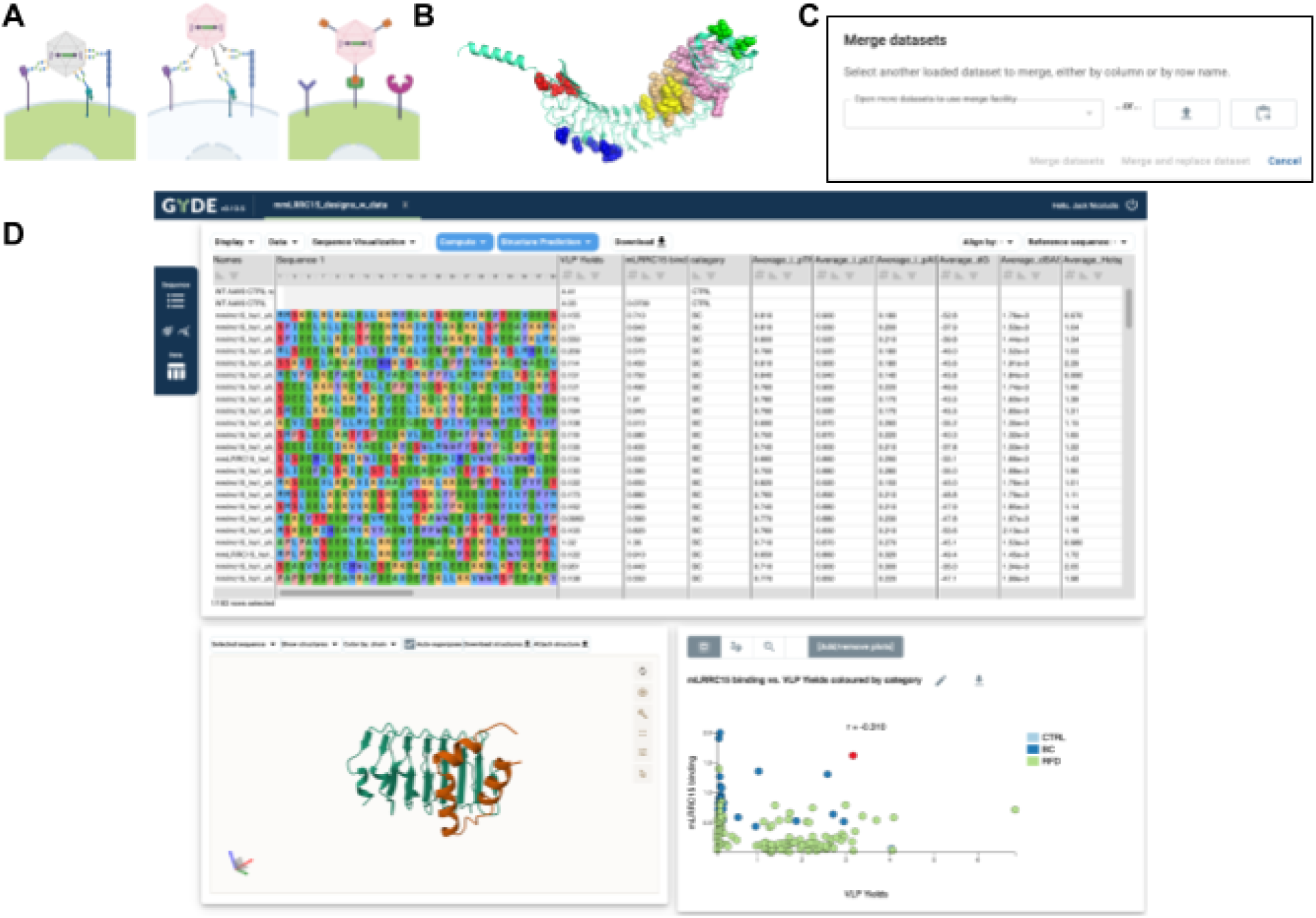
GYDE enables *de novo* binder generation (A) AAVs can be re-targeted to new receptors by eliminating existing glycan binding through mutagenesis and incorporating novel *de novo* binding motifs (in orange) into the AAV capsid protein. Image created in BioRender ^51^. (B) Hotspots were manually defined on the surface of an AlphaFold model of LRRC15. (C) Experimental data can be merged into existing GYDE session using the “Merge datasets” feature which allows adding files or copy-paste table data and merging based on desired columns. (D) The resulting GYDE session with experimental data (VLP yield and mLRRC15 binding) alongside predicted structures of the binders and a table-view of all designs enable analyzing sequence-structure-function relationships.

## Discussion

GYDE addresses a critical need in the field of protein science and drug discovery by providing an open-source, web-based collaborative platform that integrates computational analyses with intuitive visual interfaces. The platform’s core strength lies in its ability to democratize access to cutting-edge AI models and computational tools for protein and antibody structure prediction, engineering, design, and downstream analyses, making these resources easily accessible to both novice and expert protein scientists.

The GYDE architecture is characterized by its modular design, which facilitates flexibility and scalability. By integrating various components such as the GYDE UI, GYDE Server, external computational and data resources, the system offers a comprehensive platform for collaborative protein data analysis. The architecture supports efficient data handling, user interaction, and computational processing, making it well-suited for the demands of modern data-driven protein science environments. The underlying data model, based on simple data frames, enhances flexibility and ease of data manipulation and visualization. Furthermore, GYDE’s data management system, backed by MongoDB, provides robust and scalable storage while enabling real-time collaboration and seamless data sharing among researchers. By utilizing the Slivka compute API deployed on premise or at a remote site, GYDE leverages high-performance computing resources for computationally intensive tasks like AlphaFold2 and ProteinMPNN. This integration reduces the fragmentation often associated with traditional computational approaches. GYDE’s modular architecture, encompassing computation, visualization, and data management, ensures a comprehensive and seamless data analysis experience. The interface is designed to facilitate interaction with protein data through components like the MSA viewer, Mol* structural visualization, and plotting tools. The platform also supports the integration of experimental data for comparative analysis alongside computational predictions. These views comport with the well-known sequence-structure-function relationship that is central to protein science, and by fully encompassing this relationship, GYDE allows distinct view points onto key and relevant data across a broad range of techniques and algorithms in this domain.

We highlighted the utility of GYDE through several case studies spanning various protein science applications in drug discovery. These include target discovery (STM multimers), benchmarking new methods (comparative analysis of co-folding methods), antibody and protein engineering efforts (anti-PD-1 antibody, TEV protease), and de novo protein design (LRRC15). These examples demonstrate some of the breadth of applications that are possible in GYDE. GYDE is also flexibly built to incorporate new tools as they become available, ensuring it stays relevant to the quickly changing landscape of computational drug discovery.

While GYDE enables running jobs through the platform directly, large scale analyses (e.g. thousands of structure predictions) are still better suited outside the platform where compute resources can be managed more directly. However, GYDE’s efficient and flexible data import options allow facile visualization, interpretation, and sharing of results. This can also be beneficial in the evaluation and assessment stage with new algorithms before full integration. Similarly, generating conformational dynamics of proteins is not currently supported in GYDE, but we imagine that the GYDE platform interface and data model could be used to represent each row in a dataset as a distinct conformational state across a molecule dynamics trajectory.

We anticipate the effective integration of Large Language Models (LLMs) to create prompt-based access to GYDE capabilities as a replacement to point-and-click interfaces. In our experience the analysis of the output data will still need effective visualization methods, and we believe the GYDE interface, its intuitive presentation of the sequence-structure-function relationship, and its ‘send to GYDE’ (or model context protocol MCP) options for seamless connectivity would be effective ways for these multi-agentic AI systems to hybridize with GYDE.

GYDE represents a significant advancement in enabling collaborative and accessible protein science research in drug discovery. By tightly integrating computational tools, intuitive visualizations, and robust data management within a web-based platform, GYDE lowers the barrier to entry for advanced computational analyses, empowers bench scientists to independently explore complex questions, and fosters more efficient interdisciplinary collaborations. The demonstrated utility across various case studies underscores its potential to accelerate drug discovery and protein engineering efforts, making it a valuable tool for both academic and commercial research environments. The flexibility of the system and the ease with which new tools and models can be added position GYDE as a dynamic platform that can adapt to the rapidly evolving landscape of computational protein science and provide benefit as a shared community platform.

## Supporting information

GYDE Supplementary

Supp_ExAntibodyCoV2.xlsx

## Acknowledgements

We’d like to thank Tadej Satler for running the RFDiffusion binder design workflow for LRRC15 and Kapil Bajaj for running Alphafold2 at scale on HPC systems.

## Code Availability

The GYDE source repository is available at https://github.com/proteinverse/gyde.

## References

1. Jumper, J. et al. Highly accurate protein structure prediction with AlphaFold. Nature 596, 583–589 (2021).

2. Baek, M. et al. Accurate prediction of protein structures and interactions using a three-track neural network. Science 373, 871–876 (2021).

3. Ahdritz, G. et al. OpenFold: retraining AlphaFold2 yields new insights into its learning mechanisms and capacity for generalization. Nat Methods 21, 1514–1524 (2024).

4. The OpenFold3 Team. OpenFold3-preview. Zenodo 10.5281/ZENODO.17485510 (2025).

5. Wohlwend, J. et al. Boltz-1 Democratizing Biomolecular Interaction Modeling. Preprint at 10.1101/2024.11.19.624167 (2024).

6. Passaro, S. et al. Boltz-2: Towards Accurate and Efficient Binding Affinity Prediction. Preprint at 10.1101/2025.06.14.659707 (2025).

7. Chai Discovery et al. Chai-1: Decoding the molecular interactions of life. Preprint at 10.1101/2024.10.10.615955 (2024).

8. Robust deep learning–based protein sequence design using ProteinMPNN. https://www.science.org/doi/10.1126/science.add2187doi:10.1126/science.add2187.

9. Pacesa, M. et al. BindCraft: one-shot design of functional protein binders. Preprint at 10.1101/2024.09.30.615802 (2024).

10. Watson, J. L. et al. De novo design of protein structure and function with RFdiffusion. Nature 620, 1089–1100 (2023).

11. Beg, M., et al. Using Jupyter for reproducible scientific workflows. 10.48550/ARXIV.2102.09562 (2021) doi:10.48550/ARXIV.2102.09562.

12. Mirdita, M. et al. ColabFold: making protein folding accessible to all. Nat Methods 19, 679–682 (2022).

13. The Galaxy Community et al. The Galaxy platform for accessible, reproducible, and collaborative data analyses: 2024 update. Nucleic Acids Research 52, W83–W94 (2024).

14. Sayers, E. A General Introduction to the E-utilities. in Entrez Programming Utilities Help [Internet] (National Center for Biotechnology Information (US), 2022).

15. Madeira, F. et al. The EMBL-EBI Job Dispatcher sequence analysis tools framework in 2024. Nucleic Acids Research 52, W521–W525 (2024).

16. Waterhouse, A. M., Procter, J. B., Martin, D. M. A., Clamp, M. & Barton, G. J. Jalview Version 2—a multiple sequence alignment editor and analysis workbench. Bioinformatics 25, 1189–1191 (2009).

17. The PyMOL Molecular Graphics System, Version 3.0 Schrödinger, LLC. Schrodinger, LLC.

18. Anfinsen, C. B., Haber, E., Sela, M. & White, F. H. THE KINETICS OF FORMATION OF NATIVE RIBONUCLEASE DURING OXIDATION OF THE REDUCED POLYPEPTIDE CHAIN. Proc. Natl. Acad. Sci. U.S.A. 47, 1309–1314 (1961).

19. Katoh, K. & Standley, D. M. MAFFT Multiple Sequence Alignment Software Version 7: Improvements in Performance and Usability. Molecular Biology and Evolution 30, 772–780 (2013).

20. Absolve antibody variable domain sequence analysis https://github.com/Genentech/Absolve. Genentech (2025).

21. Sehnal, D. et al. Mol* Viewer: modern web app for 3D visualization and analysis of large biomolecular structures. Nucleic Acids Research 49, W431–W437 (2021).

22. Abanades, B. et al. ImmuneBuilder: Deep-Learning models for predicting the structures of immune proteins. Commun Biol 6, 575 (2023).

23. Warowny, M., MacGowan, S., Barton, G. & Procter, J. Slivka and Slivka-bio: a lightweight, unopinionated framework for executables as web services, and its application to bioinformatics. https://f1000research.com/posters/10-707 (2021) doi:10.7490/f1000research.1118680.1.

24. Molecular Operating Environment (MOE), Scientific Vector Language (SVL) source code provided by Chemical Computing Group ULC, 910-1010 Sherbrooke St. W., Montreal, QC H3A 2R7, 2025.

25. Dreyer, F. A., et al. Conformation-Aware Structure Prediction of Antigen-Recognizing Immune Proteins. Preprint at 10.48550/ARXIV.2507.09054 (2025).

26. Raybould, M. I. J., Turnbull, O. M., Suter, A., Guloglu, B. & Deane, C. M. Contextualising the developability risk of antibodies with lambda light chains using enhanced therapeutic antibody profiling. Commun Biol 7, (2024).

27. Park, E. & Izadi, S. Molecular Surface Descriptors to Predict Antibody Developability. Preprint at 10.1101/2023.07.18.549448 (2023).

28. Blaabjerg, L. M. et al. Rapid protein stability prediction using deep learning representations. eLife 12, (2023).

29. Leaver-Fay, A. et al. Rosetta3. in Methods in Enzymology vol. 487 545–574 (Elsevier, 2011).

30. Dieckhaus, H., Brocidiacono, M., Randolph, N. Z. & Kuhlman, B. Transfer learning to leverage larger datasets for improved prediction of protein stability changes. Proc. Natl. Acad. Sci. U.S.A. 121, (2024).

31. Dauparas, J. et al. Atomic context-conditioned protein sequence design using LigandMPNN. Nat Methods 22, 717–723 (2025).

32. Siepe, D. H. et al. Identification of orphan ligand-receptor relationships using a cell-based CRISPRa enrichment screening platform. eLife 11, e81398 (2022).

33. Banhos Danneskiold-Samsøe, N., et al. AlphaFold2 enables accurate deorphanization of ligands to single-pass receptors. Cell Systems 15, 1046–1060.e3 (2024).

34. Evans, R. et al. Protein complex prediction with AlphaFold-Multimer. Preprint at 10.1101/2021.10.04.463034 (2021).

35. Bryant, P., Pozzati, G. & Elofsson, A. Improved prediction of protein-protein interactions using AlphaFold2. Nat Commun 13, (2022).

36. Krissinel, E. & Henrick, K. Inference of Macromolecular Assemblies from Crystalline State. Journal of Molecular Biology 372, 774–797 (2007).

37. Szklarczyk, D. et al. STRING v11: protein–protein association networks with increased coverage, supporting functional discovery in genome-wide experimental datasets. Nucleic Acids Research 47, D607–D613 (2019).

38. Zhang, J. et al. Computing the Human Interactome. Preprint at 10.1101/2024.10.01.615885 (2024).

39. Škrinjar, P., Eberhardt, J., Durairaj, J. & Schwede, T. Have protein-ligand co-folding methods moved beyond memorisation? 2025.02.03.636309 Preprint at 10.1101/2025.02.03.636309 (2025).

40. Barnes, C. O. et al. SARS-CoV-2 neutralizing antibody structures inform therapeutic strategies. Nature 588, 682–687 (2020).

41. Sampei, Z. et al. Antibody engineering to generate SKY59, a long-acting anti-C5 recycling antibody. PLoS ONE 13, e0209509 (2018).

42. Briney, B., Inderbitzin, A., Joyce, C. & Burton, D. R. Commonality despite exceptional diversity in the baseline human antibody repertoire. Nature 566, 393–397 (2019).

43. Olsen, T. H., Boyles, F. & Deane, C. M. Observed Antibody Space: A diverse database of cleaned, annotated, and translated unpaired and paired antibody sequences. Protein Science 31, 141–146 (2022).

44. Bernardes, J. P. et al. Longitudinal Multi-omics Analyses Identify Responses of Megakaryocytes, Erythroid Cells, and Plasmablasts as Hallmarks of Severe COVID-19. Immunity 53, 1296–1314.e9 (2020).

45. Galson, J. D. et al. Deep Sequencing of B Cell Receptor Repertoires From COVID-19 Patients Reveals Strong Convergent Immune Signatures. Front. Immunol. 11, (2020).

46. Nielsen, S. C. A. et al. Human B Cell Clonal Expansion and Convergent Antibody Responses to SARS-CoV-2. Cell Host & Microbe 28, 516–525.e5 (2020).

47. Liang, W.-C. et al. Structure- and machine learning-guided engineering demonstrate that a non-canonical disulfide in an anti-PD-1 rabbit antibody does not impede antibody developability. mAbs 16, 2309685 (2024).

48. Sumida, K. H. et al. Improving Protein Expression, Stability, and Function with ProteinMPNN. J. Am. Chem. Soc. 146, 2054–2061 (2024).

49. Phan, J. et al. Structural Basis for the Substrate Specificity of Tobacco Etch Virus Protease. Journal of Biological Chemistry 277, 50564–50572 (2002).

50. Ray, U. et al. Exploiting LRRC15 as a Novel Therapeutic Target in Cancer. Cancer Research 82, 1675–1681 (2022).

51. Bainbridge, T. Created in BioRender. (2025).

