## Supplementary material for "GYDE: A collaborative drug discovery platform for AI-powered protein design and engineering": GYDE Supplementary

### Methods

**AAV engineering**

We cloned mammalian expression constructs coding for these binders as recombinant insertions into the VR4 loop of AAV9 VP3 capsid subunit, at position G455, flanked by five-residue GGAGG linkers. The VR4 loop has been previously demonstrated to accommodate larger, globular domains [1]. Virus-like particles (VLPs) were generated by co-transfecting HEK293 cells with expression plasmids coding for the modified VLPs as well as AAV9 Assembly-Activating Protein (AAP), which is required for efficient capsid assembly, at a 5:1 mass ratio, respectively. Seven days post-transfection, the conditioned media containing the VLPs was isolated. The relative yields and mmLRRC15 binding of virus-like particles (VLPs) containing the binder insertions were evaluated via biolayer interferometry (BLI), using biosensors with either immobilized mouse LRRC15 extracellular domain or immobilized AAVX, a pan-AAV binding nanobody. The yield and binding data were then merged into the GYDE session to allow inspection of the results (Figure XC). The resulting session allows a tight coupling of the design data, predicted structures and experimental results that enable a data-rich analysis environment that can be easily explored by bench scientists (Figure XD).

**Single pass membrane protein interactome**

A library of 2,523 single-pass transmembrane (STM) receptor ectodomain prey constructs and 707 human STM query proteins was generated based on computational predictions, functional annotations, and manual curation[2,3]. The receptor ectodomains were cloned into pRK5 vectors (Genentech) with a C-terminal hIgG (Fc) tag, while STM query proteins were fused to the pentameric helical region of rat cartilage oligomeric matrix protein (COMP) and β-lactamase to enable a colorimetric readout with the nitrocefin substrate. Both libraries were transiently transfected into Expi293 cells using 25 kDa linear polyethyleneimine (PEI), and conditioned media containing secreted proteins were harvested 7 days post-transfection. Receptor-ligand screening was performed using a high-throughput AVEXIS-based platform[3,4] in 384-well Protein A-coated plates, which were incubated overnight at 4°C with conditioned media containing the STM receptor library to immobilize prey proteins. Plates were washed with PBS containing $Ca^{2+}$ and $Mg^{2+}$ and subsequently incubated with conditioned media containing pentamerized STM proteins for 1 hour at room temperature. After washing to remove unbound proteins, nitrocefin substrate (Calbiochem) was added to detect β-lactamase activity, indicative of protein-protein interactions. Absorbance at 485 nm was measured using a Tecan Infinite M1000 multimode microplate reader. Interaction data were normalized by subtracting the background plate absorbance (10th percentile) and scaling to the maximal absorbance (99.5th percentile). A supervised random forest classifier was implemented using the Caret R package, trained on a benchmark dataset of validated positive and negative interactions. High-confidence receptor-ligand interactions were identified based on class probabilities: P(Positive)>=0.75, P(Non-specific) <= 0.25 and P(Negative) <= 0.05.

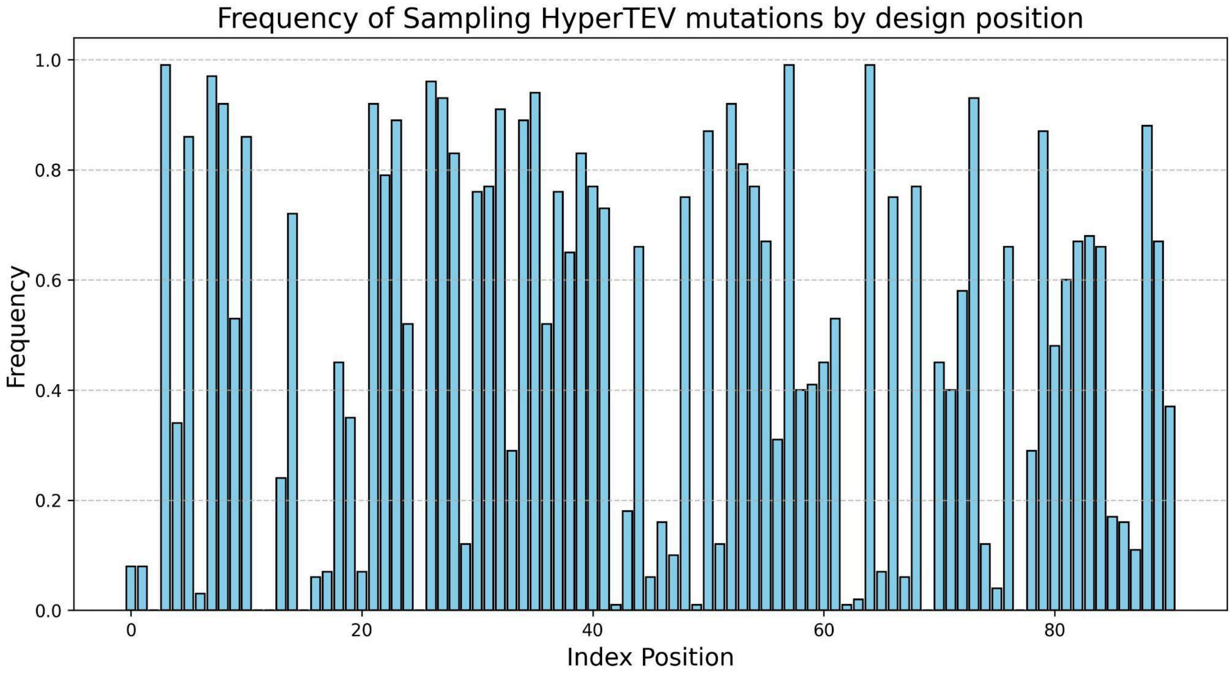

Figure S2. Histogram plot across the TEV protein sequence positions permitted to vary (x-axis) indicating our results' frequency of sampling an amino acid identical to one of the 3 experimentally verified HyperTEV designs from published studies.

### References


1. Judd, J. *et al.* Random Insertion of mCherry Into VP3 Domain of Adeno-associated Virus Yields Fluorescent Capsids With no Loss of Infectivity. *Molecular Therapy - Nucleic Acids* **1**, e54 (2012).
2. Clark, H. F. *et al.* The Secreted Protein Discovery Initiative (SPDI), a Large-Scale Effort to Identify Novel Human Secreted and Transmembrane Proteins: A Bioinformatics Assessment. *Genome Res.* **13**, 2265–2270 (2003).
3. Martinez-Martin, N. *et al.* An Unbiased Screen for Human Cytomegalovirus Identifies Neuropilin-2 as a Central Viral Receptor. *Cell* **174**, 1158-1171.e19 (2018).
4. Bushell, K. M., Söllner, C., Schuster-Boeckler, B., Bateman, A. & Wright, G. J. Large-scale screening for novel low-affinity extracellular protein interactions. *Genome Res.* **18**, 622–630 (2008).
